# Information theory rules out the reflex-chain model of *C. elegans* locomotion

**DOI:** 10.1101/2022.02.01.478702

**Authors:** John Webb, Saul Kato

**Author notes:** JW wrote the code for the project and performed the experiments. JW and SK worked together on all other aspects of the project, including conceptualization, data analysis, and writing.

## Abstract

Despite decades of research, whether the *C. elegans* traveling-wave sinusoidal body pattern during locomotion is produced (a) by the undulations of the head followed by wave propagation down the body, or (b) via centrally coordinated posture control along the body, is still under debate. By studying relationships between the time series of postural angles along the body extracted from videos of moving worms, we find that the reflex-chain model can be refuted, in both forward and backward locomotion as well as during swimming and crawling behaviors. We show that information theory applied to animal behavior can yield insights into the neural control of behavior.

---

**H**ow the nematode *C. elegans* moves in a well-executed serpentine fashion is still unknown despite a detailed anatomical knowledge, connectome and genetic access to each of its 302 neurons (1, 2). The body motor system of *C. elegans* consists of overlapping 95 body wall muscle cells that ring the body and 75 body motor neurons grouped into 12 similar neuromuscular units running down the body (3). Two main models exist for *C. elegans* locomotion: a reflex-chain model where the dorsoventral undulations of the head set up an oscillatory pattern that propagates down the body via connections between adjacent neuromuscular units and biomechanical linkage, and an alternative active posture model where the sinusoidal body posture along the entire body is effected by active neural control not solely deriving from lateral neuromuscular signaling from the head to tail (Fig 1a). The earliest computer simulations of *C. elegans* movement were based on a reflex-chain model, and more recent simulations based on proprioceptive reflex chains recapitulate aspects of *C. elegans* movement (4–7). Worms crawl on their side with a smoothly propagating sinusoidal undulation with little body slippage outside of the sinusoidal path they trace out on their crawling surface; we surmise that the appearance of a smooth and consistent traveling wave inspired the reflex-chain model.

**Fig. 1.**
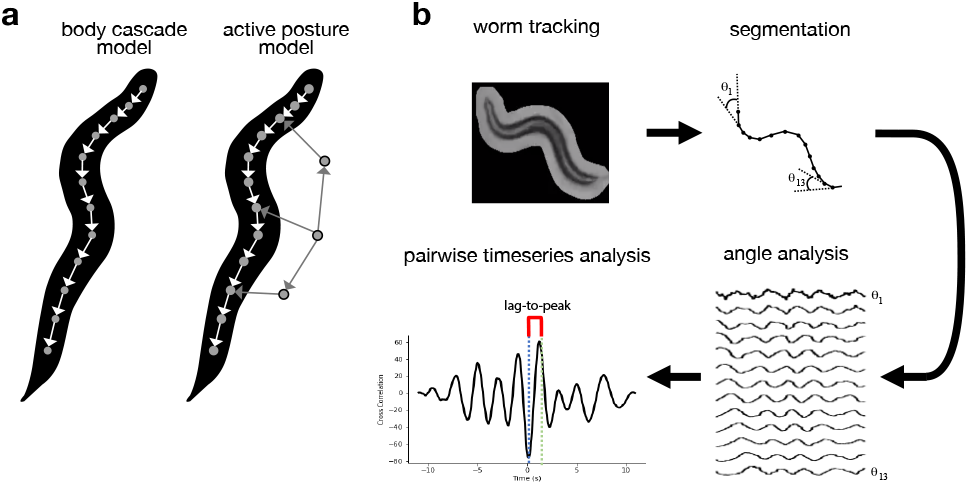
Locomotion models and analysis overview. a. Schematics of the reflex-chain model (left) versus the active posture model (right). b. Overview of analysis of worm movement. Sustained bouts of forward locomotion were tracked, worm skeletons were segmented, and the worm tangent angles were plotted versus time for each segment. Finally, pairwise measures – time lag to peak cross-correlation, peak absolute cross-correlation, and mutual information were calculated for all joint angle pairs.

An alternative model, which we term the active posture model, posits that worm motion is driven by multiple sites of centrally coordinated neural signals along the body. (Fig. 1a). These signals may be produced by a pattern generator (CPG) consisting of one more cells. Recently, rhythmically active groups of neurons for forward and backward locomotion have been identified (8–12), but whether these groups of neurons represent autonomous CPGs is still to be resolved.

## Results

### Cross-correlation of postural angle time series reveals non– monotonic noise accumulation down the body

To generate quantitative worm movement data, we recorded high resolution videos of worms crawling on an agar surface using a custom-built motorized-stage microscope and image–based tracking software system (13) and performed video analysis (14) to silhouette and segment the worms. We then extracted time series of the postural tangent angles between each of 13 body segments (Fig. 1b). As expected, these time series resembled a series of phase-lagged noisy sinusoids. Performing analysis with a finer discretization of body segments did not change the key findings.

The sinusoidal appearance of the signals suggested that cross-correlation analysis would be revealing. The cross-correlation of two closely related sinusoidal signals in the presence of noise consists of a set of peaks of decaying magnitude (Fig. 1b). The x-coordinate of the peak of the cross-correlation provides an estimate of the time lag of the signals. The maximum absolute value of the cross-correlation provides a scalar estimate of the relatedness of the signals measured at the most favorable relative time delay, and it is reduced by the amount of noise present in the transformation between the signals. To simulate the undulations of forward locomotion under the reflex-chain model, we created a sine wave to represent head postural angle time series, added noise and a phase delay to the signal to generate the posteriorly adjacent postural angle time series, and iterated this procedure down the body.

We computed the cross-correlation between each body joint angle with respect to the anterior-most (head) joint angle during this simulated pattern of forward locomotion (Fig. 1b). As expected, in the reflex-chain model simulation, the time lag to peak cross-correlation with respect to the first segment time series increased monotonically, and the peak absolute cross-correlation with respect to the first segment time series decreased monotonically with increasing segment number (Fig. 2a,b). We then performed the same analysis of our experimental data. For this analysis, we selected contiguous time series sections when the animal was crawling forward and not turning. In our experimental worms, we did not observe a stably increasing time lag to peak cross-correlation (Fig. 2a), and strikingly, we observed a strong breaking of monotonicity in the peak absolute cross-correlation (Fig. 2b). There were local minima in the peak absolute cross-correlation of angle pairs (1,5) and (1,9). This deviation from monotonicity suggests that the reflex-chain model is a poor fit to experiment. However, there was trial-to-trial variability in the pattern of peak correlations and time lag; thus, we sought a more robust measure of information transmission.

**Fig. 2.**
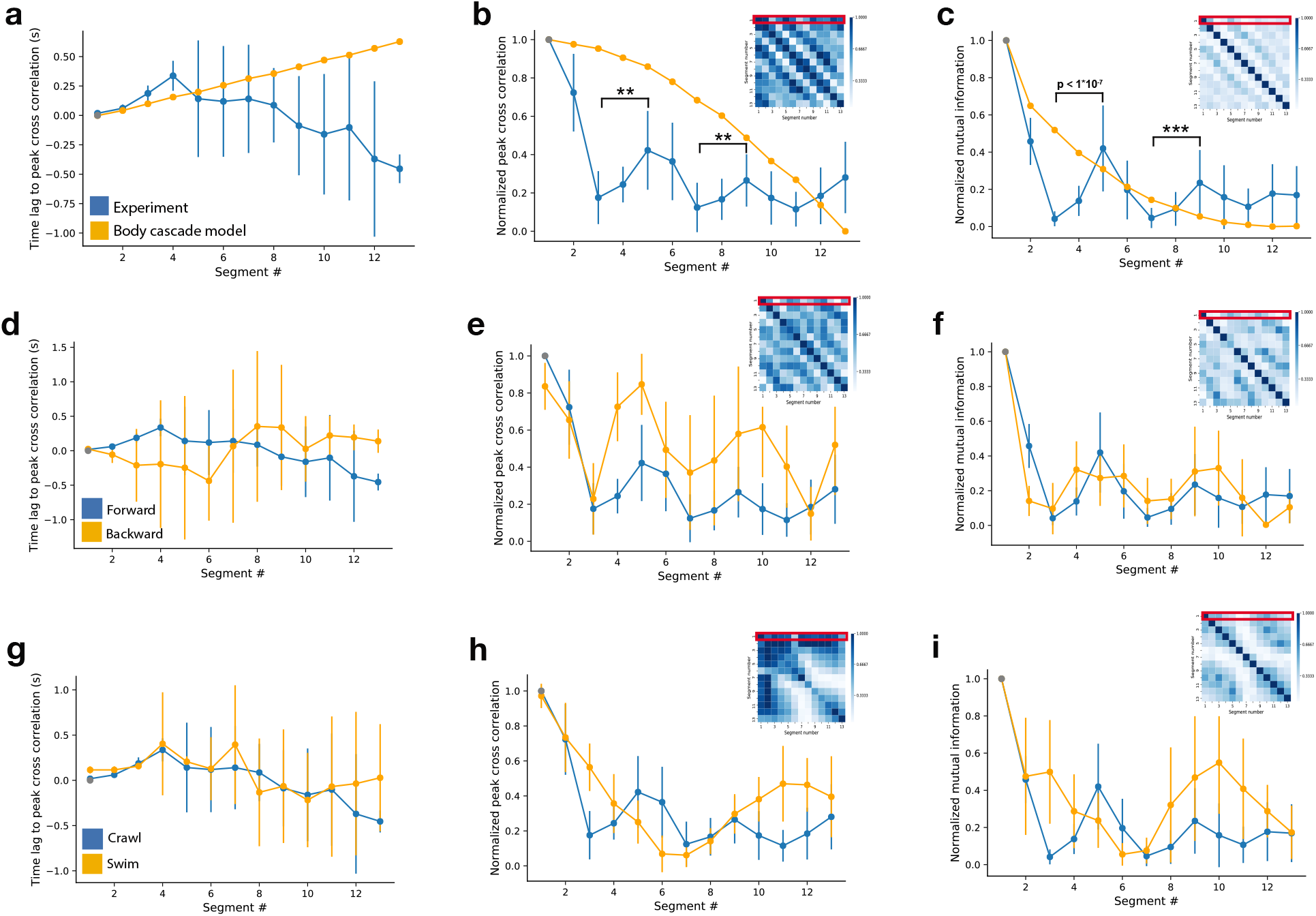
a-c: Postural angle time series relationships during forward locomotion. a. Time lag to the peak cross correlation vs angle # calculated for the reflex-chain model (orange) and wild-type worms (blue) for forward locomotion, n=10. b. Heat map shows peak abs. cross-correlations computed between all angle pairs; first row is shown in the larger plot. Note that the experimental data is non-monotonic, in contrast to the model. (Segment 5 *−* segment 3 and segment 9 *−* segment 7, random sampling with replacement, **p<0.01) c. Mutual information plotted between all angles in the inset heatmap, with the mutual information relative to the head angle plotted. (Segment 5 *−* segment 3 and segment 9 *−* segment 7, random sampling with replacement, ***p<0.001) **d-f: Forward versus backward crawling**. d. Time lag to peak cross-correlation for forward (blue) and backward (orange) locomotion (compared to the head angle for forward and tail angle for backward), n=9 worms. e. Peak abs. cross-correlation normalized to the head angle for forward locomotion (blue) and tail angle for backward locomotion (orange). f. The peak mutual information normalized to the head angle for forward (blue) and tail angle segment for backward (orange). **g-i: Forward crawling versus swimming**. g. Time lag to peak cross-correlation for crawling (blue) and swimming (orange) locomotion compared to the anterior-most segment. n=10 worms for each group. h. Peak abs. cross-correlation for crawling (blue) and swimming (orange) locomotion normalized to the anterior-most segment. i. The mutual information for crawling (blue) and swimming (orange) locomotion normalized to the anterior-most segment.

### Mutual information suggests centrally coordinated posture control

A central theorem of information theory is the *data processing inequality*: a propagating signal can only lose, and not gain information from transmission from point to point, due to the accumulation of noise (15). If a worm moved according to the reflex-chain model, the mutual information between the head joint angle and each successive body angle would monotonically decrease (Fig. 2b, c). However, we found a strong experimental deviation from monotonic information loss. The two local maxima of the mutual information relative to angle 1 occur at the same angle numbers (5 and 9) as the two local maxima of the peak absolute cross-correlation, suggesting that active postural control may be transmitted to the periphery through two specific points. We also measured the mutual information between all angle pairs (Fig. 2c, inset).

### Forward crawling, backward crawling, and swimming are under centrally coordinated postural control

We extended our analysis to backwards locomotion, in this case using the posterior-most (tail) angle as angle 1. *C. elegans* backwards locomotion is shorter in bout duration and crawling length than forwards locomotion, so we employed shorter time windows than those used for forward locomotion. Similar to forward locomotion, we found a non-monotonic peak absolute cross-correlation and non-monotonic mutual information (Fig. 2e, f). The peak absolute cross-correlation has local maxima for angle pairs (1,5) and (1,10) (Fig. 2e, f). A reflex-chain model can thus be rejected for both directions of crawling, and both appear to coordinate control at two points along the body.

We then analyzed worm swimming. It has been argued that *C. elegans* swimming and crawling represent distinct neural control patterns (16) rather than solely the result of biomechanical influence of a changing physical substrate. We found the time-lag to absolute cross-correlation to be non-monotonic but, in contrast to the crawling state, the peak absolute cross-correlation has only one, rather than two local maxima, suggesting a different mode of central cooordination. (Fig. 2g, h). The reflex-chain model can be rejected for swimming worms as well as crawling worms.

## Discussion

We claim that the reflex-chain model of worm movement is inconsistent with fine analyses of behavioral data. Our data suggests there are two body locations where central coordination reaches the periphery. With higher resolution video recordings, detailed anatomical registration of neural and neuron-to-muscle connectivity data from the worm connectome could suggest particular neurons and connections responsible for centrally coordinated posture control.

Our data is consistent with recent loss-of-function studies. One study showed that forward-rhythm undulations persist in posterior body segments even when anterior body segments are paralyzed (8). Another study found that when anterior A motor neurons were ablated, it did not prevent the propagation of reversal waves in posterior body segments (9). In addition to recent studies suggesting the presence of neural oscillators, there is also evidence for lateral information transmission between adjacent neuromuscular units (17). If there are multiple CPG groups driving locomotion, our data suggest that they are strongly coupled. We hypothesize that coordinated oscillatory postural control signals reach the neuromuscular periphery at two specific points along the body, bypassing intervening neuromuscular units. These signals are shaped into a spatially smooth traveling body waveform by lateral neuromuscular signal transmission and further smoothed by biomechanical linkage.

We assume that there is not severe segment-to-segment heterogeneity in the noise accumulated during the local biomechanical transformation from muscle to body bend angle; if this transformation noise were both strong and wildly different along the body, it could undermine our interpretation of the non-monotonicity of our measures. However, we find this unlikely given the robustness of the results and lack of an intuition as to how such heterogeneity might occur.

## Materials and Methods

We recorded videos of wild-type (N2) worms using a custom tracking microscope and TierpsyTracker software (13, 14). We manually identified bouts of forward crawling, backward crawling, and swimming. Analysis code is available at https://github.com/focolab/worm-locomotion-control and was written in python.

## ACKNOWLEDGMENTS

SK is funded by the NIGMS (R35GM124735), the Alfred Sloan Foundation, and the Weill Neurohub. The CGC is funded by NIH Office of Research Infrastructure Programs (P40 OD010440).

